# New evidence of a rhythmic priming effect that enhances grammaticality judgments in children

**DOI:** 10.1101/193961

**Authors:** Alexander Chern, Barbara Tillmann, Chloe Vaughan, Reyna L. Gordon

## Abstract

Musical rhythm and the grammatical structure of language share a surprising number of characteristics that may be intrinsically related in child development. The present study aimed to understand the potential influence of musical rhythmic priming on subsequent spoken grammar task performance in children with typical development who were native speakers of English. Participants (ages 5 to 8 years) listened to rhythmically regular and irregular musical sequences (within-subjects design) followed by blocks of grammatically correct and incorrect sentences upon which they were asked to perform a grammaticality judgment task. Rhythmically regular musical sequences improved performance in grammaticality judgment compared to rhythmically irregular musical sequences. No such effect of rhythmic priming was found in two non-linguistic control tasks, suggesting a neural overlap between rhythm processing and mechanisms recruited during grammar processing. These findings build on previous research investigating the effect of rhythmic priming by extending the paradigm to a different language, testing a younger population, and employing non-language control tasks. These findings of an immediate influence of rhythm on grammar states (temporarily augmented grammaticality judgment performance) also converge with previous findings of associations between rhythm and grammar traits (stable, generalized grammar abilities) in children. Taken together, this study provides additional evidence for shared neural processing for language and music, and warrants future investigations of potentially beneficial effects of innovative musical material on language processing.

## Introduction

Framed by a large literature showing associations between language and music skills in children, there is great interest in the possibility that one domain may impact the other via shared neural resources (Kraus & Slater, 2016). Sensitivity to musical features in particular has been hypothesized to be fundamental to early language and grammar acquisition (Brandt, Gebrian, & Slevc, 2012). Grammar, also called morpho-syntax, is the use of rules about how words change their form and combine with other words to make phrases and sentences that enable individuals to communicate with each other. Recent evidence suggests that children with enhanced rhythm perception ability tend to have better spoken grammar skills (Gordon, Shivers, et al., 2015) and children with grammatical deficits tend to have impaired rhythm (Cumming, Wilson, Leong, Colling, & Goswami, 2015). This shared variance could be explained in part by similarities in how rhythm and grammar employ hierarchical structures emerging from rule-based expectancies that unfold over time at multiple levels (Fitch & Martins, 2014) and more generally, by shared brain mechanisms for temporal attention (Jones & Boltz, 1989).

Several studies have explored potential shared neural resources for rhythm/timing and syntactic processing in adults, using the event-related potential (ERP) method. For instance, altering temporal intervals between word onsets has shown that regular, predictable presentation improves syntactic processing, reflected by an increase of the P600 ERP component (Schmidt-Kassow & Kotz, 2009). Interestingly, prior listening to rhythmically regular musical stimuli restored the (otherwise missing) P600 to grammatical (linguistic) violations in patients with basal ganglia lesions (Kotz, Gunter, & Wonneberger, 2005) and Parkinson’s disease (Kotz & Gunter, 2015), thus suggesting that rhythmic stimulation can improve detection of grammatical violations in subsequently presented sentences.

To evaluate potential benefits of rhythmic listening on subsequent syntactic processing in children with typical development (TD) and their peers struggling with language, Przybylski et al. (2013) tested native French-speaking children (ages 6 to 11 years) with TD, specific language impairment, and dyslexia. Performance on a grammaticality judgment task improved after listening to a rhythmically regular musical sequence (characterized by its strong metrical structure) compared to a rhythmically irregular (non-metrical) musical sequence. Converging with data demonstrating relationships between rhythm and grammar traits in children (Gordon, Jacobs, Schuele, & McAuley, 2015; Gordon, Shivers, et al., 2015), these studies show that musical rhythm with a strong beat structure can influence language states (see also, Bedoin et al., 2017; Bedoin, Brisseau, Molinier, Roch, & Tillmann, 2016).

Our aim was to test this *Rhythmic Priming Effect* (RPE), the positive influence of rhythmically regular musical stimulation on grammar task performance, in TD English-speaking children. Additionally, we sought to determine whether RPE (originally demonstrated in children with a mean age of 9) would be present in a younger population (mean age 6) who are still developing their grammatical skills. These investigations are needed as a future step toward use of RPE in Anglophone children in applied (educational and clinical) settings, complementing findings described above in children who speak French (a syllable-timed language; Lee & Todd, 2004, c.f., Arvaniti, 2009). Given reports of RPE in German in adult clinical populations (Kotz et al., 2005; Kotz & Gunter, 2015), we hypothesized that RPE would also be present in English-speaking children (both German and English are stress-timed languages).

Moreover, additional work on RPE included a condition with neutral environmental sounds (Bedoin et al., 2016), demonstrating that RPE is linked to a benefit from listening to regular stimuli rather than a detriment from irregular stimuli. Given that music listening might lead to small and temporary enhancements in cognitive performance (i.e., mixed findings for the Mozart effect, Pietschnig, Voracek, & Formann, 2010), it is also crucial to test whether RPE could be attributed to a general effect of arousal on task performance or a more specific effect from shared neural mechanisms between rhythm and grammar processing. We thus tested the influence of rhythmic priming on two non-linguistic (math and visuospatial) control tasks.

## Methods

### Participants

Sixteen native English-speaking children aged 5;6 to 8;7 (M = 6;5, SD = 11 months; 9 males, 7 females) participated in this study, which received Vanderbilt University Institutional Review Board approval. Parents gave informed consent; children gave verbal assent. Parents reported no concerns about cognitive, motor, emotional, sensory, or language development. Participants were recruited from a larger ongoing study examining rhythm and language, and all scored greater than 85 on all subtests of the Test of Language Development (Newcomer & Hammill, 2008) with non-verbal IQ in normal range (M = 117.4, SD = 18.6) on the Primary Test of Nonverbal Intelligence (Ehrler & McGhee, 2008). Hearing was screened and all participants were found to have normal hearing. Parent education level (proxy for socioeconomic status) and music experience as assessed in Gordon, Shivers, et al. (2015) are reported and described in Supplementary Table 1, along with other participant characteristics. Children received a small toy, and parents received a $30 gift card for compensation. Study data were managed and stored using REDCap electronic data capture tools (Harris et al., 2009).

### Rhythmic priming effect paradigm – grammaticality judgment task

Musical stimuli. Stimuli consisted of the same two 32-s musical sequences from Przybylski et al. (2013) (Supplementary Fig. 1), containing same number of tones, but differing in rhythmic structure such that meter was strong (regular) or weak/unmetrical (irregular).

Linguistic Stimuli. Material was composed of 72 grammatically correct (36) and violation (36) English sentences. Each grammatically correct sentence had an equivalent violation sentence; each sentence was heard once in either the correct or incorrect form. Two types of violations were used: subject-verb agreement (e.g. *Every day, the mom toast/toasts the bread for their breakfast*) or tense agreement (e.g. *Yesterday, the men punched/punch a bag at the gym*). Sentences were recorded by a female speaker in an anechoic chamber and edited in PRAAT software (Boersma, 2001) with a cross-splicing method (from onset of the verb clause), allowing each violation sentence to be created from two grammatically correct sentences (e.g., underlined portion of *<u>Yesterday, the men</u> punched a bag at the gym,* was combined with underlined portion of *Yesterday, the men <u>punch a bag at the gym</u>*), ensuring they were spoken with natural prosody. Sentences had an average length of 9.72 words (range = 9–10) and 13.22 syllables (range = 10–16) and average duration of 3044 ms (±498). Two experimental lists were generated such that sentences were counterbalanced across participants and conditions (regular vs. irregular primes and correct vs. incorrect grammar).

Experimental paradigm. Each block of sentences contained three grammatically correct and three violation sentences, with a total of 36 sentences across the six blocks (randomized within each block). Half the blocks were preceded by the regular stimulus and half by the irregular stimulus (see Fig. 1; pairing of prime and sentence blocks were counterbalanced across participants; primes alternated between regular and irregular).

**Fig. 1.**
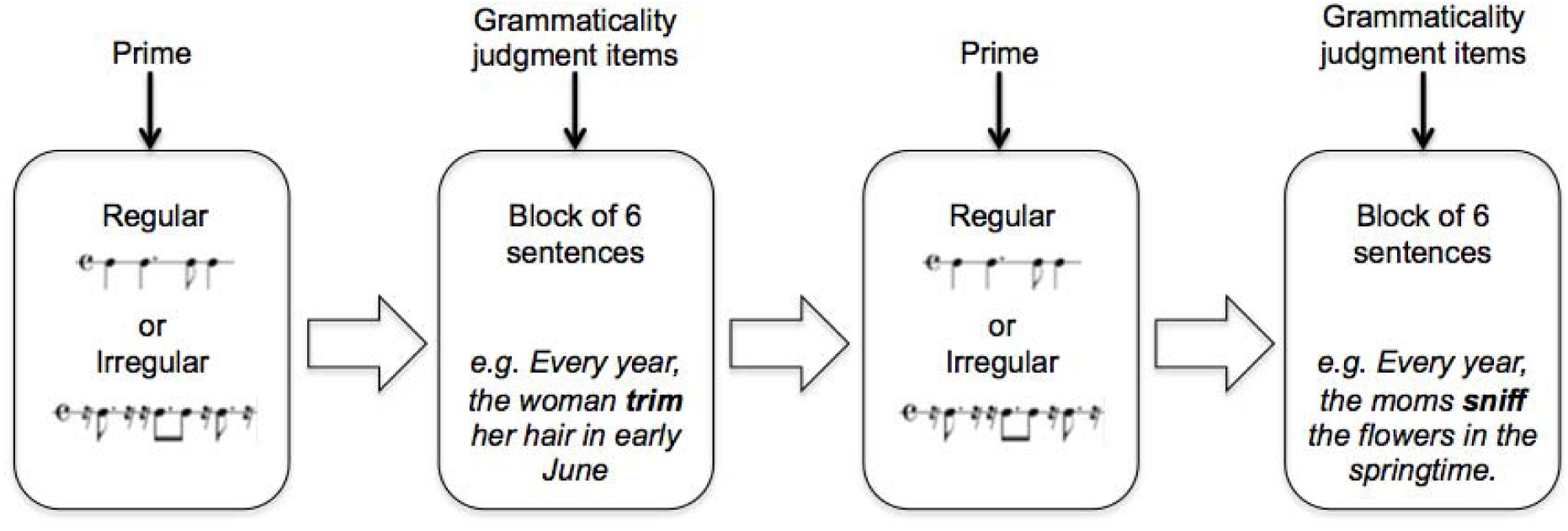
Rhythmic Priming Effect Experimental Paradigm. The musical prime (regular or irregular) is heard once, and then a block of 6 sentences is presented. Children are asked to perform grammaticality judgment on each sentence item.

### Procedure

Testing was presented as a computer game. For each block, children were asked to listen to the music while being shown a guitar symbol on the screen. A sentence was then presented auditorily, and children were asked to indicate whether the “correct” or “incorrect” dragon had spoken by pointing to the appropriate picture (dragons appear on the screen on opposite sides). Before starting, participants were administered four practice items (two grammatically correct and two violations) without music and explained that “this dragon (satisfied-looking dragon) always says things right, and this dragon (confused-looking dragon) always says things wrong.” If participants provided the wrong answer for any practice items, the experimenter explained why the sentence was correct or incorrect. Sounds were presented over speakers (Alesis Elevate 5) hand-calibrated to 70 dB at a range of 3 feet.

**Control tasks.** Two non-linguistic control tests were utilized with the rhythmic priming paradigm. The *math task* consisted of two forms (19 items each; equivalent in content and difficulty) derived from the Wide Range Achievement Test 4 (Wilkinson & Robertson, 2006), and utilized a mix of visual and auditory items to assess counting, identifying numbers, and solving simple problems (e.g. *“How many are 3 apples and 4 apples?”)*. Each form was preceded by either the regular or irregular rhythmic sequence. In both control tasks, order of musical sequences and forms was counterbalanced across participants. Forms for each condition were scored as proportion of total items answered correctly. The *visuo-spatial task* consisted of the cancellation subtest from the Wechsler Intelligence Scale for Children, fourth edition (Wechsler, 2003). As with the math task, two equivalent forms (32 items each) were used with each participant, each preceded by a musical sequence. Children were asked to mark animal targets interspersed among common non-animal targets (e.g. car, tree) within 23 seconds. Forms were scored as total number of targets marked correctly subtracted by number of targets marked incorrectly. All participants completed the math task, and N=11 completed the cancellation task (due to experimenter logistics).

### Data Analysis

Grammaticality judgment performance was subjected to signal detection analysis, yielding *d’*, a measure of discriminability, and *c,* a measure of response bias (Macmillan & Creelman, 2005). Paired t-tests were performed to determine if there was significant difference in performance after listening to regular stimuli compared to irregular stimuli. To estimate effect sizes, we calculated Cohen’s *d* (small = 0.20, medium = 0.5, or large = 0.8).

**Fig. 2.**
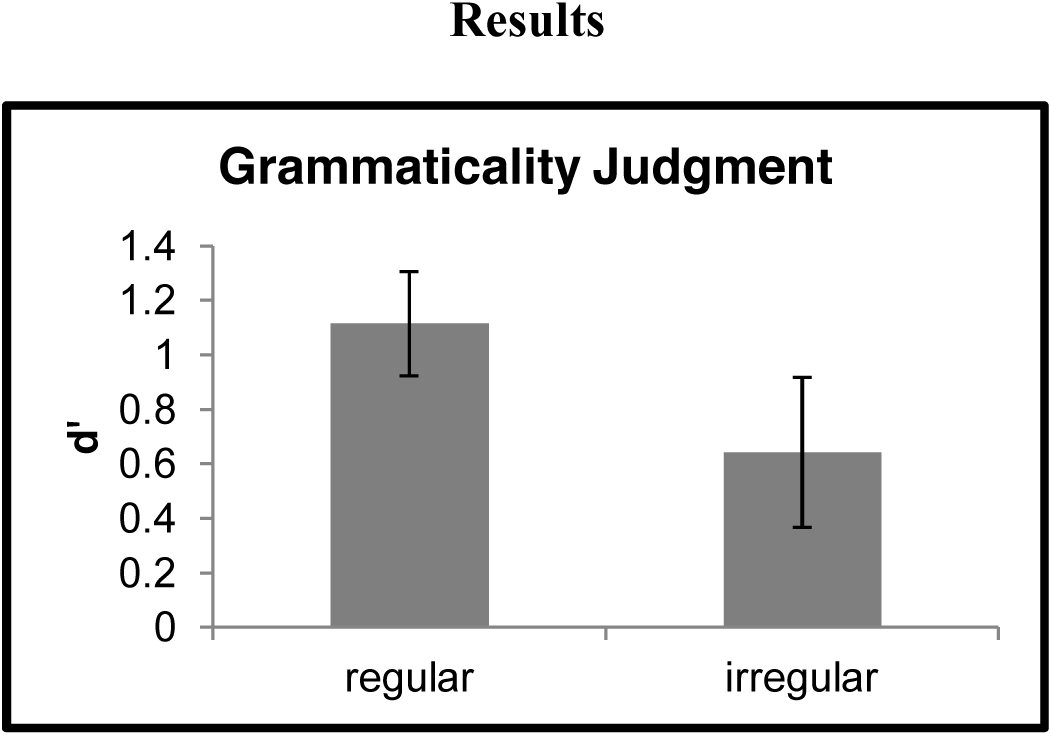
Rhythmic priming effect on grammaticality judgment task. Findings in N=16 children show that sentences preceded by regular primes led to better grammaticality judgment performance.

Grammaticality judgment performance was significantly better (t = 3.07, df = 15, *p* = 0.008, Cohen’s *d* = 0.57, medium-sized effect; see Fig. 2) after listening to regular musical sequence primes (mean *d’ = 1.11*, SE = 0.18) compared to irregular musical sequence primes (mean *d’* = 0.60, SE = 0.26). Performance after listening to regular or irregular stimuli did not change significantly between the first and second half of each block (Supplementary Table 2, Supplementary Fig. 2).

Analysis of *c* showed relatively small response biases. Participants showed a slight bias to respond “incorrect” after having listened to regular musical stimuli (*c* = −0.22±0.08) and irregular musical stimuli (*c* = −0.04±0.12). There was no significant difference in *c* for the two conditions (t = −1.72, df = 15, *p =* 0.11, Cohen’s *d* = 0.45).

After listening to regular vs. irregular stimuli (mean_reg_ and mean_irreg_, respectively), there were no significant differences in visuo-spatial test performance (*p* = 0.63; mean_reg_ correct = 15.90, SE = 0.99; mean_irreg_ correct = 15.60, SE = 1.26) or math test performance (*p =* 0.15; mean_reg_ proportion correct = 0.81, SE = 0.04; mean_irreg_ proportion correct = 0.84, SE = 0.04) (Supplementary Table 3).

## Discussion

This study examined whether rhythmically regular and irregular musical stimuli differentially influenced performance of typically developing (TD), English-speaking children in a subsequent grammar task. As hypothesized, performance improved after listening to regular musical stimuli compared to irregular stimuli. This effect of the musical prime on the d’ of the grammar task was of medium effect size. However, for the non-linguistic control tasks there was no significant difference in math or visuo-spatial performance after listening to regular versus irregular musical stimuli. Thus, the findings in the language task likely cannot be attributed to general cognitive benefits due to enhanced arousal from listening to musical stimuli (e.g., Pietschnig et al., 2010) and rather suggest a more specific sharing of neural resources between rhythm and grammar. In comparison to other work showing that use of rhythmic primes matched to the exact prosodic features of subsequently presented speech can enhance speech processing (Cason & Schon, 2012), our study showed the influence of musical stimuli with a rhythmic structure that did not match the speech rhythm of our sentences. Our finding converges with recent work in French-speaking children (Przybylski et al., 2013), suggesting RPE not only occurs in French (and German; Kotz et al., 2005), but also English and potentially other languages. Moreover, here we observed RPE in a younger population compared to prior work, highlighting possible relevance for language development. Intriguingly, in this study and prior work, RPE occurs with enhanced grammar task performance in children with typical language (and thus no deficit in grammatical skill), suggesting a robust benefit of rhythmic listening.

Adding to the growing literature showing shared processing of music and linguistic stimuli (e.g., Kraus & Slater, 2016), our results are consistent with theories of overlapping early development of rhythm and language (Brandt et al., 2012). The process of prosodic bootstrapping suggests that toddlers make use of prosodic cues, such as timing and intonation of phrase boundaries, to parse syntax during language acquisition (de Carvalho, Dautriche, Lin, & Christophe, 2017). Sensitivity and preference to sounds that fit the patterns of native meter in both the musical and linguistic domains may develop in parallel (Jusczyk, 2000). Meter perception from rhythmic patterns has been demonstrated in infants with a variety of methods (e.g. Hannon & Johnson, 2005; Zhao & Kuhl, 2016), suggesting that the distinction between regular and irregular stimuli employed by RPE is highly relevant to the development of music perception from a very early age.

Sensitivity to rhythmic structures of music and speech continues to play an important role in normal and disordered language acquisition (Cumming et al., 2015). The present findings converge with recent work showing benefits from short-term passive musical listening on grammar processing in children with disordered language (Bedoin et al., 2016; Przybylski et al., 2013). Taken together, these results suggest an immediate activation of shared or connected brain networks, involving even less effort than that required by music training (see Patel, 2014 and Fujii & Wan, 2014).

A potential mechanism driving RPE can be framed in the context of *Dynamic Attending Theory*, which hypothesizes that rhythmic patterns in speech and music enables entrainment of attention and generates temporal expectancies for future events, facilitating perception, segmentation, and integration (Jones & Boltz, 1989). Such entrainment has been suggested to occur through neural resonance (Large, Herrera, & Velasco, 2015), enabling hierarchical organization of stimuli with different subdivisions. As proposed in prior work, rhythmic priming may modulate temporal attention, mobilizing neural oscillators also involved in generating hierarchical expectancies that allow listeners to parse the speech stream and process syntax (Przybylski et al., 2013; Bedoin et al., 2016).

Neural mechanisms supporting RPE could include the basal ganglia, which play a role in both musical beat perception and language processing (Krishnan, Watkins, & Bishop, 2016; Merchant, Grahn, Trainor, Rohrmeier, & Fitch, 2015); neural responses to grammatical violations by rhythmic priming were restored in individuals with basal ganglia lesions (Kotz & Gunter, 2015; Kotz et al., 2005). Left inferior frontal gyrus (LIFG) plays an important role in hierarchical processing of language and music (Fitch & Martins, 2014) and may also be involved in brain networks that lead to RPE. Moreover, imaging evidence in jazz musicians indicates that detection of deviations in musical rhythm are associated with activation of areas involved in linguistic syntax, including LIFG (Donnay, Rankin, Lopez-Gonzalez, Jiradejvong, & Limb, 2014). Further work is needed to understand the neural processes and regions driving RPE and to what degree generalized dynamic attending processes are recruited. Kotz, Schwartze, and Schmidt-Kassow (2009) proposed two pathways (pre-supplementary motor area (SMA)-basal ganglia, and cerebellar-thalamic-pre-SMA) involved in sequencing and temporal attention. Potential clinical benefits of RPE in individuals with atypical language, who may show deficits in one pathway, may arise from rhythmic compensation through the other pathway or enhanced use of the impaired one (Przybylski et. al, 2013).

To have clinical relevance, therapeutic interventions employing this rhythmic priming must induce a stable trait effect on language ability, as opposed to a temporary, state effect. It will be important to quantify how many sessions (perhaps combining RPE and language treatment activities or active rhythm tasks) are needed to reach sustained improvements in grammatical skill (Schon & Tillmann, 2015). This prospect is especially promising for remediating the reported co-morbid grammatical and rhythm deficits in children with language impairments (Cumming et al., 2015) and for use during language therapy sessions in children with cochlear implants (Bedoin et al., 2017).

Future work should also address some of the limitations of the current study. First, the oral grammaticality judgment task, while relevant for school activities, is not as ecological as a conversational language task would be. Investigations of RPE on spoken elicitation tasks should be thus considered (e.g., Hidalgo, Falk, & Schon, 2017), and additional non-linguistic control conditions with more trials could be interspersed within the same protocol. Moreover, study of this phenomenon in larger samples will allow for the exploration of individual differences in temporal processing ability and language skill as predictors of benefits derived from RPE (e.g., associations between rhythm and grammar skills, Gordon, Shivers, et al., 2015), and for the pursuit of mediating neural mechanisms. More broadly, this line of research could address questions about whether the rhythmic structure of background music could affect our efficacy of reading, writing, and speaking.

In conclusion, there is an accumulation of evidence for a positive impact of listening to regular musical rhythms (with a strong metrical structure) on subsequent grammatical (syntactic) processing. This RPE has been observed in school-aged children with typical and atypical language development, in adults with basal ganglia lesions and Parkinson’s disease, in native speakers of French and German, and is due to a benefit of regular musical rhythms rather than a detriment of irregular rhythms (Bedoin et al., 2016; Kotz & Gunter, 2015; Kotz et al., 2005; Przybylski et al., 2013). The present study enhances this literature by demonstrating RPE in English, in younger children, and that RPE is a shared effect on rhythm and grammar rather than a more general effect of music on cognition. Taken together with recent reports of individual differences in musical rhythm perception that predict grammar traits (Gordon, Shivers, et al., 2015), these findings suggest rhythmic stimuli (i.e. music) can influence language processing states (i.e., grammar task performance).

## Acknowledgments

The authors thank Allison Aaron, Katherine Margulis, Kristin Gummersheimer, and CJ Walters for assistance in data collection and entry, Sonja Kotz, Eleanor Harding, Rachel Waters, and Carolyn Shivers for collaboration on sentence stimuli on a prior unpublished study, and Miriam Lense for insight on the experimental paradigm. This research was supported by funding from Vanderbilt Medical Scholars, NIH R03DC014802, and Vanderbilt Trans-Institutional Programs. Use of REDCap was made possible by Vanderbilt CTSA grant UL1 TR000445 from NCATS/NIH.

## References

Arvaniti, A. (2009). Rhythm, timing and the timing of rhythm. Phonetica, 66(1-2), 46–63.

Bedoin, N., Besombes, A. M., Escande, E., Dumont, A., Lalitte, P., & Tillmann, B. (2017). Boosting syntax training with temporally regular musical primes in children with cochlear implants. Ann Phys Rehabil Med.

Bedoin, N., Brisseau, L., Molinier, P., Roch, D., & Tillmann, B. (2016). Temporally Regular Musical Primes Facilitate Subsequent Syntax Processing in Children with Specific Language Impairment. Front Neurosci, 10, 245.

Boersma, P. (2001). Praat, a system for doing phonetics by computer. Glot International, 5(9/10), 341–345.

Brandt, A., Gebrian, M., & Slevc, L. R. (2012). Music and early language acquisition. Front Psychol, 3, 327.

Cason, N., & Schon, D. (2012). Rhythmic priming enhances the phonological processing of speech. Neuropsychologia, 50(11), 2652–2658.

Cumming, R., Wilson, A., Leong, V., Colling, L. J., & Goswami, U. (2015). Awareness of Rhythm Patterns in Speech and Music in Children with Specific Language Impairments. Front Hum Neurosci, 9, 672.

de Carvalho, A., Dautriche, I., Lin, I., & Christophe, A. (2017). Phrasal prosody constrains syntactic analysis in toddlers. Cognition, 163, 67–79.

Donnay, G. F., Rankin, S. K., Lopez-Gonzalez, M., Jiradejvong, P., & Limb, C. J. (2014). Neural substrates of interactive musical improvisation: an FMRI study of ‘trading fours’ in jazz. PLoS One, 9(2), e88665.

Ehrler, D. J., & McGhee, R. L. (2008). PTONI: primary test of nonverbal intelligence: Pro-Ed.

Fitch, W. T., & Martins, M. D. (2014). Hierarchical processing in music, language, and action: Lashley revisited. Ann N Y Acad Sci, 1316, 87–104.

Fujii, S., & Wan, C. Y. (2014). The Role of Rhythm in Speech and Language Rehabilitation: The SEP Hypothesis. Front Hum Neurosci, 8, 777.

Gordon, R. L., Jacobs, M. S., Schuele, C. M., & McAuley, J. D. (2015). Perspectives on the rhythm-grammar link and its implications for typical and atypical language development. Ann N Y Acad Sci, 1337, 16–25.

Gordon, R. L., Shivers, C. M., Wieland, E. A., Kotz, S. A., Yoder, P. J., & Devin McAuley, J. (2015). Musical rhythm discrimination explains individual differences in grammar skills in children. Dev Sci, 18(4), 635–644.

Hannon, E. E., & Johnson, S. P. (2005). Infants use meter to categorize rhythms and melodies: implications for musical structure learning. Cogn Psychol, 50(4), 354–377.

Harris, P. A., Taylor, R., Thielke, R., Payne, J., Gonzalez, N., & Conde, J. G. (2009). Research electronic data capture (REDCap)-a metadata-driven methodology and workflow process for providing translational research informatics support. J Biomed Inform, 42(2), 377–381.

Hidalgo, C., Falk, S., & Schon, D. (2017). Speak on time! Effects of a musical rhythmic training on children with hearing loss. Hear Res, 351, 11–18.

Jones, M. R., & Boltz, M. (1989). Dynamic attending and responses to time. Psychol Rev, 96(3), 459–491.

Jusczyk, P. W. (2000). The discovery of spoken language (1st MIT Press pbk. ed.). Cambridge, Mass.: MIT Press.

Kotz, S. A., & Gunter, T. C. (2015). Can rhythmic auditory cuing remediate language-related deficits in Parkinson’s disease? Ann N Y Acad Sci, 1337, 62–68.

Kotz, S. A., Gunter, T. C., & Wonneberger, S. (2005). The basal ganglia are receptive to rhythmic compensation during auditory syntactic processing: ERP patient data.

Kotz, S. A., Schwartze, M., & Schmidt-Kassow, M. (2009). Non-motor basal ganglia functions: a review and proposal for a model of sensory predictability in auditory language perception. Cortex, 45(8), 982–990.

Kraus, N., & Slater, J. (2016). Beyond Words: How Humans Communicate Through Sound.Annu Rev Psychol, 67, 83–103.

Krishnan, S., Watkins, K. E., & Bishop, D. V. (2016). Neurobiological Basis of Language Learning Difficulties. Trends Cogn Sci, 20(9), 701–714.

Large, E. W., Herrera, J. A., & Velasco, M. J. (2015). Neural Networks for Beat Perception in Musical Rhythm. Front Syst Neurosci, 9, 159.

Lee, C. S., & Todd, N. P. (2004). Towards an auditory account of speech rhythm: application of a model of the auditory ‘primal sketch’ to two multi-language corpora. Cognition, 93(3), 225–254.

Macmillan, N. A., & Creelman, C. D. (2005). Detection Theory: A User’s Guide: Lawrence Erlbaum Associates.

Merchant, H., Grahn, J., Trainor, L., Rohrmeier, M., & Fitch, W. T. (2015). Finding the beat: a neural perspective across humans and non-human primates. Philos Trans R Soc Lond B Biol Sci, 370(1664), 20140093.

Newcomer, P. L., & Hammill, D. D. (2008). TOLD-P:4: Test of Language Development. Austin: Pro-Ed.

Patel, A. D. (2014). Can nonlinguistic musical training change the way the brain processes speech? The expanded OPERA hypothesis. Hear Res, 308, 98–108.

Pietschnig, J., Voracek, M., & Formann, A. K. (2010). Mozart effect–Shmozart effect: A meta-analysis. Intelligence, 38(3), 314–323.

Przybylski, L., Bedoin, N., Krifi-Papoz, S., Herbillon, V., Roch, D., Léculier, L., … Tillmann, B. (2013). Rhythmic auditory stimulation influences syntactic processing in children with developmental language disorders. Neuropsychology, 27(1), 121–131.

Schmidt-Kassow, M., & Kotz, S. A. (2009). Event-related brain potentials suggest a late interaction of meter and syntax in the P600. J Cogn Neurosci, 21(9), 1693–1708.

Schon, D., & Tillmann, B. (2015). Short- and long-term rhythmic interventions: perspectives for language rehabilitation. Ann N Y Acad Sci, 1337, 32–39.

Wechsler, D. (2003). WISC-IV: Administration and Scoring Manual: Harcourt Assessment.

Wilkinson, G. S., & Robertson, G. J. (2006). WRAT 4: wide range achievement test; professional manual. Lutz, FL: Psychological Assessment Resources, Inc.

Zhao, T. C., & Kuhl, P. K. (2016). Musical intervention enhances infants’ neural processing of temporal structure in music and speech. Proc Natl Acad Sci U S A, 113(19), 5212–5217.

